# The ‘hidden side’ of spin labeled oligonucleotides: Molecular Dynamics study focusing on the EPR-silent components of base pairing

**DOI:** 10.1101/537324

**Authors:** Sarath Chandra Dantu, Giuseppe Sicoli

**Affiliations:** Theoretical & Computational Biophysics Department, Max Planck Institute for Biophysical Chemistry, Am Faßberg 11, 37077, Go◻ttingen, Germany; Department of Computer Science–Synthetic Biology Theme, Brunel University London, Uxbridge, United Kingdom; Research group Electron Paramagnetic Resonance, Max Planck Institute for Biophysical Chemistry, Am Faßberg 11, 37077, Go◻ttingen, Germany; Laboratoire Avancé de Spectroscopie pour les Interactions, la Réactivité et l’Environnement (LASIRE), CNRS Lille, UMR 8516, Bâtiment C4 – Université de Lille, Sciences et Technologies, Avenue Paul Langevin 59655 Villeneuve-d’Ascq Cedex, France

## Abstract

Nitroxide labels are combined with nucleic acid structures and studied using electron paramagnetic resonance experiments (EPR). As X-ray/NMR structures are unavailable with the nitroxide labels, detailed residue level information, down to atomic resolution, about the effect of these nitroxide labels on local RNA structures is currently lacking. This information is critical to evaluate the choice of spin label. In this study, we compare and contrast the effect of TEMPO-based (N^T^) and rigid spin (Ç) labels (in both 2’-O methylated and not-methylated forms) on RNA duplexes. We also investigate sequence-dependent effects of N^T^ label on RNA duplex along with the more complex G-quadruplex RNA. Distances measured from molecular dynamics simulations between the two spin labels are in agreement with the EPR experimental data. To understand the effect of labeled oligonucleotides on the structure, we studied the local base pair geometries and global structure in comparison with the unlabeled structures. Based on the structural analysis, we can conclude that TEMPO-based and Ç labels do not significantly perturb the base pair arrangements of the native oligonucleotide. When experimental structures for the spin labelled DNA/RNA molecules are not available, general framework offered by the current study can be used to provide information critical to the choice of spin labels to facilitate future EPR studies.

**Graphical abstract:** 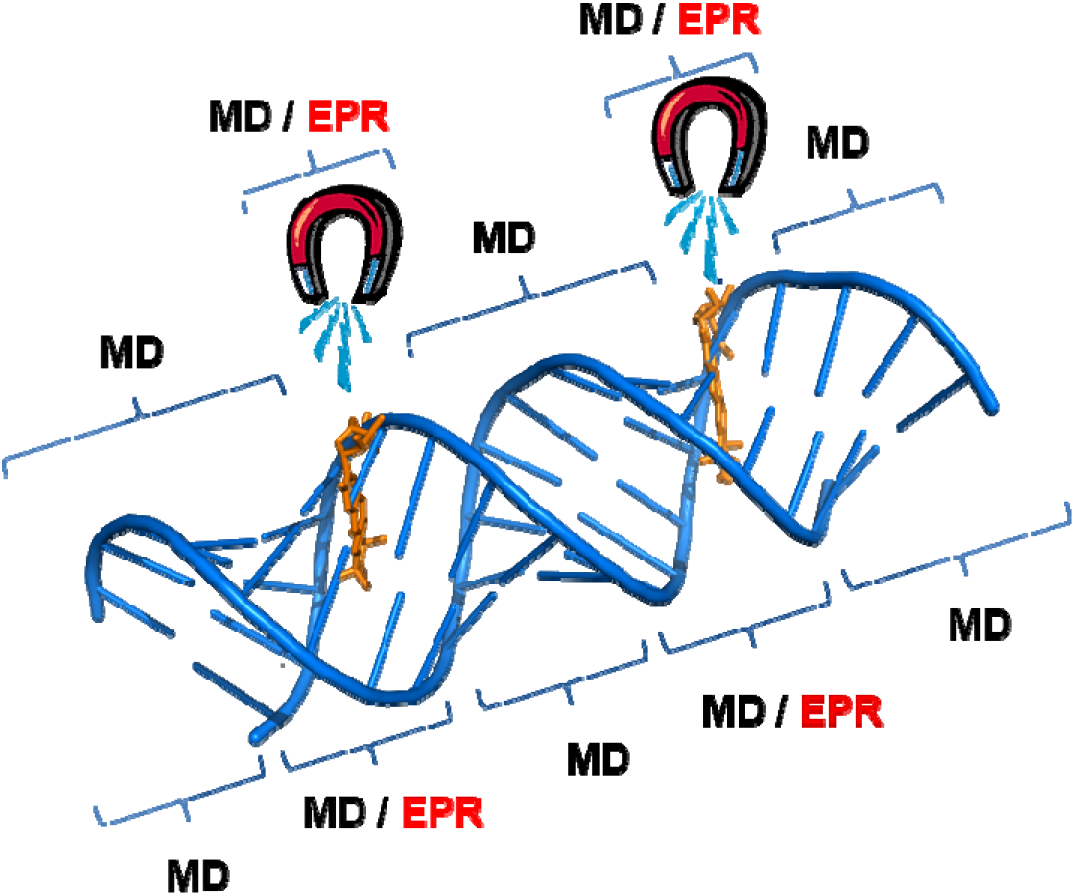

## 1. INTRODUCTION

Determining long-range inter-spin distances and their distributions from Electron Paramagnetic Resonance (EPR) experiments represents a valuable tool to elucidate the structural features of proteins and nucleic acids.[1,2] In the last few years remarkable efforts have been made towards EPR method developments and analysis.[3,4] Furthermore, advanced studies also focused on the elucidation of the relative orientation between the spin-probes.[5] A combined strategy of Molecular dynamics (MD) simulations and EPR experiments has been extensively used to determine the structural properties of deoxyribonucleic acid (DNA),[6] ribonucleic acid (RNA),[7] proteins[8] protein-RNA complexes.[9] Among the different spin probes, stable organic nitroxide radicals are widely used and in the last years their combination with metals and lanthanides rendered their applicability even more attractive.[10]

The key hypothesis for the use of such spin probes is that the Watson-Crick (WC) base pairing is preserved.[7] Under this assumption, the inter-spin distance can be determined with high accuracy. EPR studies have shown that the nitroxide labels may affect the native structure of biomolecules.[11] Further, the agreement of inter-spin distances from MD and PELDOR/DEER experiments has been extensively demonstrated, but a detailed understanding of the effect of the nitroxide labels on RNA/DNA structure is still missing.[6,12–15] Thus, a complete description of the spin-probe environment is lacking and in particular, two key-issues remain unsolved: (*i*) how does the introduction of spin-probe effect the WC base pairing (i.e. on the adjacent nucleotides to that one containing the spin probe) and (*ii*) how such perturbations can propagate to the rest of the structure.

Structural effects at residue level upon spin-labeled mutagenesis are not available from EPR experiments (i.e. the ‘hidden side’). Therefore, it is mandatory to refine and integrate spectroscopic studies with complementary techniques. This will allow us to quantitatively estimate the structural perturbation upon the insertion of new spin-probes and will enable us to point out when those perturbations are negligible, increasing the specificity of the spin-probe with respect to a DNA/RNA sequence under investigation. Therefore, we want to emphasize the need to understand the influence of these nitroxide-type labels on the structures at atomic scale.

In this work, we performed extensive MD simulations of RNA structures with either N^T^ (N = nucleotide (A, G, C) with TEMPO-based spin probe) or Ç labels[16] along with the unlabeled RNA structures. A comprehensive comparison between the native structures and the analogues derived by site-directed spin labeling techniques will highlight atomic level description on regions mainly affected by structural changes upon insertion of the label. In addition to the comparison of inter-spin distance from MD simulations with the experimental EPR data, we also present the propagation of local structural perturbations induced by rigid (Ç) and flexible (N^T^) spin probes with respect to the native oligonucleotides. We have focused our MD analysis on the six spin probes depicted in **Figure 1A**, attached to both duplexes and quadruplex, with the aim of providing an atomic-scale description of the spin environment and the dynamics of RNA.

**Figure 1.**
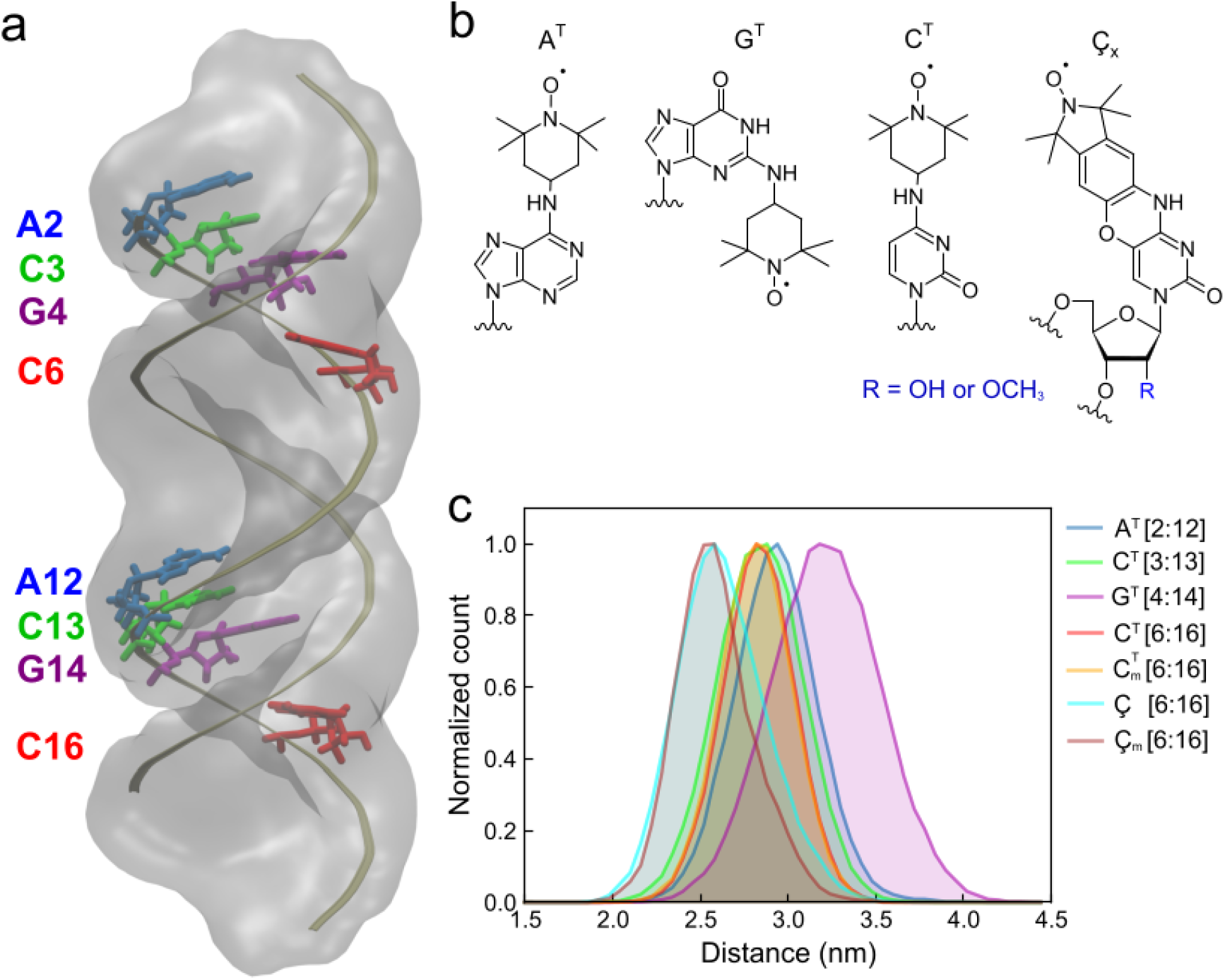
**a)** RNA duplex used in this study. Nucleic backbone is shown in gold and the eight nucleotides to which the TEMPO or Ç labels were attached are shown in licorice. **b)** Structures of spin probes analysed in this work; **c**) Distribution of distances between the oxygen (r_OO:_ N**O**·/N**O**·) radicals in the TEMPO (N^T^) and Ç labeled RNA duplexes from molecular dynamics simulations.

## 2. METHODS

### MD Simulation setup

For the MD simulations, starting structures of 20bp duplex RNA (**Figure 1a**, **Table 1**) were created using the make nucleic acid server (http://structure.usc.edu/make-na/server.html). X-ray structure 2KBP[17] was used as the starting structure for the MD simulations of G-quadruplex RNA (QRNA). All the MD simulations were performed with GROMACS 4.5[18] package. Amber99sb forcefield with parambsc0 corrections was used for all structures.[19] Forcefield parameters for N^T^ were obtained from16 and for Ç labels (**Figure 1b**) were created using same protocol described in Stendardo et al.[20] N^T^ or Ç labels were attached at positions (**Figure 1a**) described in **Table 1** to create labeled RNA and QRNA structures from unlabeled structures (RNA_U_: unlabeled RNA duplex and QRNA_U_: unlabeled QRNA). In the EPR experiments, 2’-oxygen of cytosine was methylated; therefore, to study the effect of this methylation, we also methylated the 2’-oxygen. Forcefield for the modified cytosine was created using Gaussian03 package using the RESP protocol.[21,22] Forcefield parameters of all the modified bases are available in supplementary information.

**Table 1:**
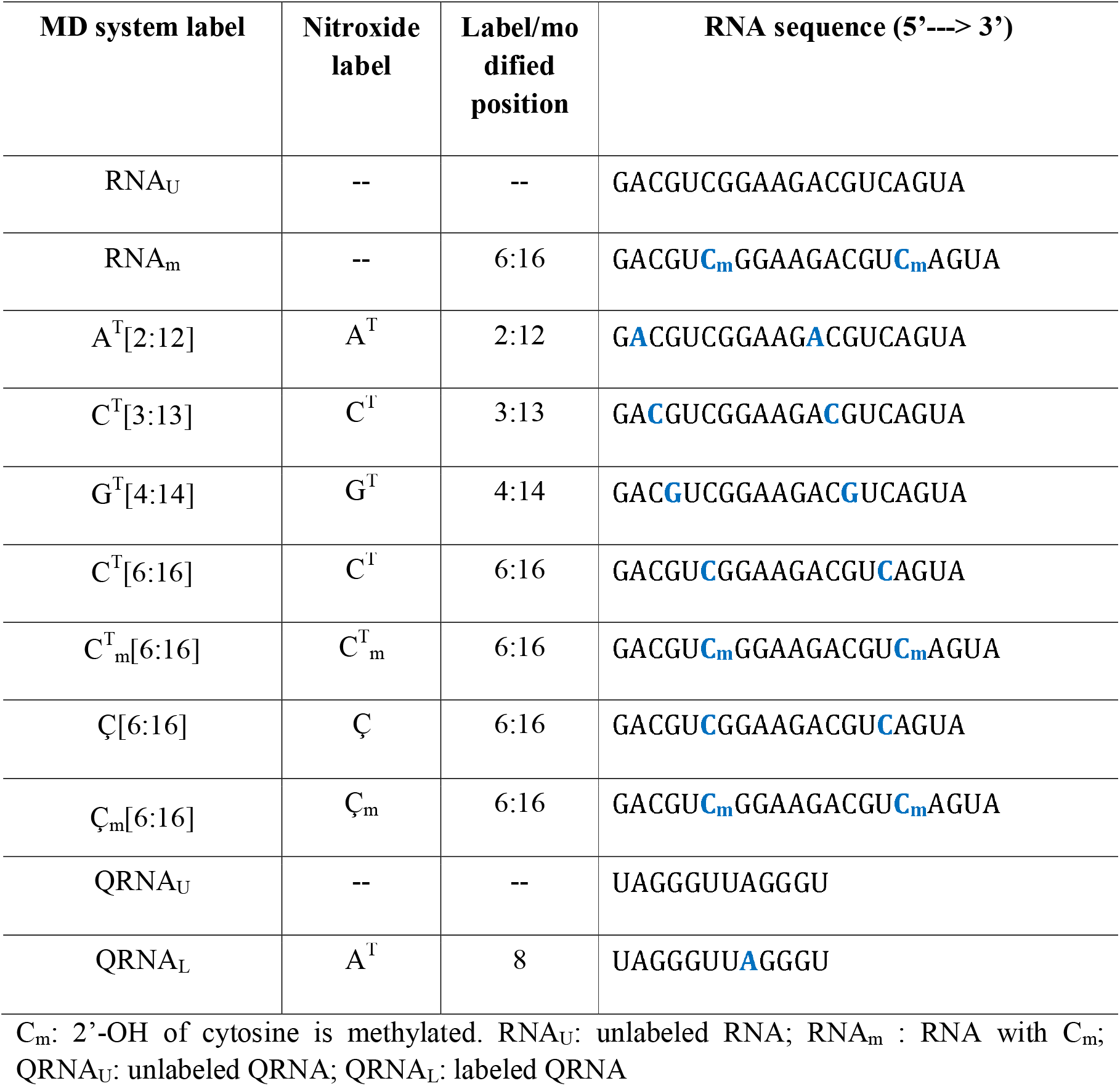
RNA sequences used for molecular dynamics simulations. Attachment site of TEMPO (N^T^) or Ç label or modified nucleotide is shown in blue. Among the eleven structures, there are nine RNA duplexes and two Quadraplex RNA (QRNA) molecules.

Each RNA molecule was placed in a dodecahedron box with a minimum distance of 1.5 nm between the molecule and the box walls. Simulation box was solvated with TIP3P18 water model. As used in the EPR experiment, 150 mM NaCl was added to the simulation box for RNA simulations. For QRNA simulations, 70 mM KCl was used. Energy minimization was performed until the largest force acting on the system was smaller than 1000 kJ mol^−1^ nm^−1^. Structure from energy minimization was equilibrated to 298K using Berendsen thermostat[23] in 100 ps using a coupling time of 0.2 ps. After temperature equilibration, using Berendsen barostat[23] and Berendsen thermostat[23] system was equilibrated to 1atm pressure in 1ns using coupling times of 1 ps and 0.2 ps respectively. The end structure of pressure equilibration was used for 110 ns production run from which the structural data for analysis was collected every 20ps. Production run simulations were performed at 298 K and 1 atm pressure by coupling the system to Nosé-Hoover thermostat[24,25] and Parrinello-Rahman barostat[26] using coupling times of 1 ps and 2 ps respectively. LINCS[27] algorithm was used to constrain bonds. Electrostatic interactions were treated by Particle-mesh Ewald algorithm[28,29] with a real space cut-off of 1.0 nm and a grid spacing of 0.12nm. Van der Waals interactions were calculated with a cut-off of 1.4 nm. Three simulations each with unlabeled RNA and QRNA and 10 simulations each for labeled structures lasting 110 ns each were performed. EPR experiments have been recorded using the PELDOR/DEER technique; such double resonance technique is based on the dipolar interaction (point-dipolar approximation) between the two spin probes. The modulation of the signal generates an oscillating signal and its frequency is proportional to the inter-spin distance.[7,30,31]

Because the flexibility of termini is much higher, the first and last two base pairs were excluded from all analysis. Nucleotide base pair geometry for RNA duplexes was analyzed using do_x3dna package[32] in three parts: inter-base pair geometry, intra-base pair geometry, and axial base pair geometry. Intra-base pair geometry along the plane of the nucleotide base pairs is defined by three *rotational* (Buckle, Propeller and Opening) and three *translational* parameters (Shear, Stretch and Stagger). Inter-base pair geometry is described by three *translational* parameters (Tilt, Roll, and Twist) and three *rotational* parameters (Rise, Shift, and Slide). Axial base pair geometry includes two *rotational* (Inclination and Tip) and two *translational* (X-displacement and Y-displacement) parameters. Thus, a set of sixteen parameters was used to systematically investigate the effect of labeling on the structure and dynamics of RNA sequences. For each base pair *i*, we calculated the average of each base pair parameter (*J*) and then with respect to the same *i^th^* residue in the unlabeled structure we calculated the difference (Δ*J_i_*). If there is no perturbation between the labeled and unlabeled base pair geometry the difference (Δ*J_i_*) would be zero:

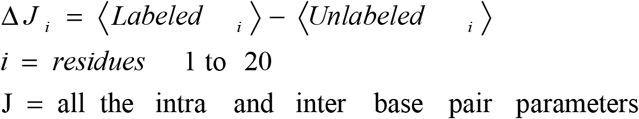

QRNA geometry parameters (area, twist, and rise) from MD trajectories were extracted using x3dna-dssr[33] and were analyzed using in-house python script. Pymol v1.8[34] and VMD v1.9[35] were used for structural visualization and analysis.

## 3. RESULTS AND DISCUSSION

We simulated a total of nine RNA duplexes and two QRNA molecules (eleven systems) with two different spin labels (**Figure 1a** **and** **1b**). In RNA duplexes the two attachment sites are 10 bp apart (2.8 nm) and 3.6 nm in QRNA. Two RNA systems (C^T^_m_ [6:16] and Ç_m_ [6:16]) have methylated cytosine at 2’-OH to study the effects of methylation on the structure (**Table 1**). For comparison purpose, unlabeled RNA duplex (RNA_U_), unlabeled RNA duplex with 2’-O methylated cytosines at 6^th^ and 16^th^ positions (RNA_m_), and unlabeled QRNA (QRNA_U_) were also simulated. We provide a comparison of inter-spin distances calculated from MD simulations and EPR data (**Table 2**) followed by a more extensive description of structural stability and geometry of base pairing. Synthesis of spin labeled sequences of oligonucleotides, for the EPR experiments, have been described elsewhere.[7,30] Pulse EPR experiments have been published elsewhere and represent the starting experimental data set. [7,30]

**Table 2:** Distances measured between the, nitrogen atoms of the native nucleobase (*r*_NL_: NL/NL), nitroxide nitrogen atoms (*r*_NN_: **N** O·/**N** O·), and the nitroxide oxygen atoms (*r*_OO_: N**O**·/N**O**·) of the two TEMPO or Çm labeled RNA, DNA, and QRNA structures in the MD simulations along with the distances measured from the EPR experiments for the respective systems.

| Label | EPR experiment | Unlabeled<br>RNA/QRNA/<br>Methylated RNA<br>($r_{NL}$) | Labeled structure | | |
| --- | --- | --- | --- | --- | --- |
| | | | Linker<br>nitrogen<br>($r_{NL}$) | Radical<br>nitrogen<br>($r_{NN}$) | Radical<br>oxygen<br>($r_{OO}$) |
| A<sup>T</sup>[2:12] | 3.1±0.2[7] | 2.7±0.1 | 2.8±0.2 | 2.9±0.2 | 3.0±0.2 |
| C<sup>T</sup>[3:13] | 3.1±0.2[7] | 2.7±0.1 | 2.8±0.2 | 2.8±0.2 | 2.9±0.2 |
| G<sup>T</sup>[4:14] | 3.2±0.3[7] | 2.8±0.2 | 3.0±0.3 | 3.2±0.3 | 3.2±0.3 |
| C<sup>T</sup>[6:16] | 3.1±0.2[7] | 2.7±0.1 | 2.8±0.2 | 2.8±0.2 | 2.9±0.2 |
| C<sup>T</sup><sub>m</sub>[6:16] | 3.1±0.2[7] | 2.7±0.2 | 2.8±0.2 | 2.8±0.2 | 2.9±0.2 |
| Ç[6:16] | n.a. | 2.7±0.2 | 2.7±0.2 | 2.6±0.2 | 2.6±0.2 |
| Ç<sub>m</sub>[6:16] | 3.1±0.3[31] | 2.7±0.2 | 2.7±0.2 | 2.7±0.2 | 2.7±0.2 |
| QRNA<sub>L</sub> | 3.77;<br>3.18* [7] | 3.5±0.5 | 3.1±0.3 | 3.7±0.3 | 3.8±0.4 |

\*Two main distances from Tikhonov regularization; error estimation as HWHH (half-width-at-half-height); The distances from MD simulations are presented with standard deviations. C<sub>m</sub>/C<sup>T</sup><sub>m</sub>: 2'-OH of cytosine is methylated. QRNA<sub>L</sub>: labeled QRNA; n.a.: not available.

### 3.1 MD distance distributions

We have monitored the distance between the nitroxide oxygen atoms (*r*_OO_: N**O**·/N**O**·) as well as the nitroxide nitrogen atoms (*r*_NN_: **N**O·/**N**O·) of the two TEMPO or Ç labels in the MD simulations. Additionally, the nitrogen atoms of the native nucleobase (*r*_NL_: NL/NL) to which the nitroxide label is attached, was used to monitor the distance between the nucleotide molecules. Since the distance between the two attachment sites cannot be measured in an EPR experiment in the unlabeled RNA, such distance for the “attachment sites” will provide an estimation of how the nitroxide inter-spin distances differ from the native structure.

Distances for the above-mentioned sequences are summarized in **Table 2** and the distance distributions obtained by DEER/PELDOR experiments are summarized in **Supporting Figure S1**. r_NN_ and r_OO_ distances, except for the Ç [6:16] and Ç_m_[6:16], are either in agreement with the EPR experimental values (G^T^[4:14]) or are within the error limit of the EPR experiments. The difference between the *r*_NN_ and *r*_OO_ distances is very small (0.1 nm) for all the duplexes. *r*_NL_ distances calculated from the unlabeled structures are shorter by 0.1 nm to 0.4 nm when compared with the *r*_NN_ and *r*_OO_ calculated from labeled structures and from the EPR data. It must be noted that distances between Ç labels (*r*_NN_ and *r*_OO_) are similar to the distances calculated between the label attachment sites in the unlabeled RNA (*r*_NL_) and 0.4 nm shorter than the distances from EPR experiments. For rest of the RNA duplexes (except G^T^[4:14], which is an exact match to EPR data), *r*_NN_ and *r*_OO_ distances from MD simulations, though shorter (0.1 nm to 0.2 nm), are within the experimental error range. [7,30] Similar trend of shorter distances between the nitroxide labels from MD simulations has been reported in previous MD studies.[6,12–15] A systemic study into the effects of forcefield parameterization of EPR labels on inter-spin distances in MD simulations are required to understand the shorter distances from MD studies.

### 3.2 Structural stability

To monitor the effect of N^T^/Ç labels on the entire structure, Root mean square deviation (RMSD) of the backbone of the entire structure (excluding the two terminal residues at 5’ and 3’ ends) with respect to the starting structure of the simulation was calculated (**Table 3**). First, we compared the effect of the methyl group at the 2’ position of the ribose sugar to unmodified RNA. The difference in backbone RMSD’s between the unlabeled RNA_m_ and unlabeled RNA is very small (0.04 nm) suggesting that methylation at 2’-oxygen have a minor effect on the RNA structure and is in agreement with earlier reported experimental and computational works.[36,37] N^T^ and Ç labeled RNA structures have higher RMSD (0.31 nm-0.34 nm) compared to the unlabeled RNA (0.24 nm). It must be noted that among the N^T^ labeled RNA duplexes, G^T^[4:14] had the lowest RMSD. The slightly higher RMSD (difference of 0.06nm to 0.1nm) of N^T^ or Ç labeled structures suggests that the effect of labels is minimal on overall structure.

**Table 3:** Root mean square deviation (RMSD) using only the phosphate backbone atoms of the entire duplex molecule excluding the two terminal residues at both 5’ and 3’ ends. Standard deviation is 0.1 nm.

| Simulation label | RMSD (nm) |
| --- | --- |
| RNA | 0.24 |
| RNA<sub>m</sub> [6:16] | 0.28 |
| A<sup>T</sup>[2:12] | 0.34 |
| C<sup>T</sup>[3:13] | 0.34 |
| G<sup>T</sup>[4:14] | 0.31 |
| C<sup>T</sup>[6:16] | 0.34 |
| C<sup>T</sup><sub>m</sub>[6:16] | 0.34 |
| Ç[6:16] | 0.33 |
| Ç<sub>m</sub>[6:16] | 0.34 |

C<sub>m</sub>/C<sup>T</sup><sub>m</sub>: 2'-OH of cytosine is methylated.

### 3.3 Base pair geometry analysis

For a more in-depth structural evaluation of the effect of nitroxide labels on the local environment at the label attachment site, we analyzed the local base pair geometry using multiple base pair parameters proposed by Dickerson et al.[38] using do_x3dna tool.[32]

#### 3.3.1 N^T^ *vs.* Ç

In C^T^[6:16] and Ç[6:16] there is no change in translational parameters when compared to the unlabeled RNA (**Supplementary Figure S2**) and perturbation trends are identical for twist, tip, and propeller (**Supplementary Figure S3**). Major differences are seen in opening, buckle, and inclination (**Figure 2**). While introduction of Ç label increases opening at the label attachment site and the preceding base pair, C^T^ label results in decreased buckling at the attachment site and decreased inclination preceding the attachment site from 6^th^ base pair and this effect is less pronounced at the internal attachment (i.e. the 16^th^ base pair). These three parameters suggest that higher ‘opening’ of the base pair is produced by Ç (increased hindrance with respect to C^T^), while in the case of ‘inclination’ there is the opposite scenario. For the changes in buckle-angle, no significant differences between the two spin probes were observed. According to these results, the choice of one spin probe with respect to the other cannot be motivated by the induced structural perturbations. However, the ease of synthesis and accessibility should also be taken into consideration, which clearly favors C^T^ over Ç.

**Figure 2.**
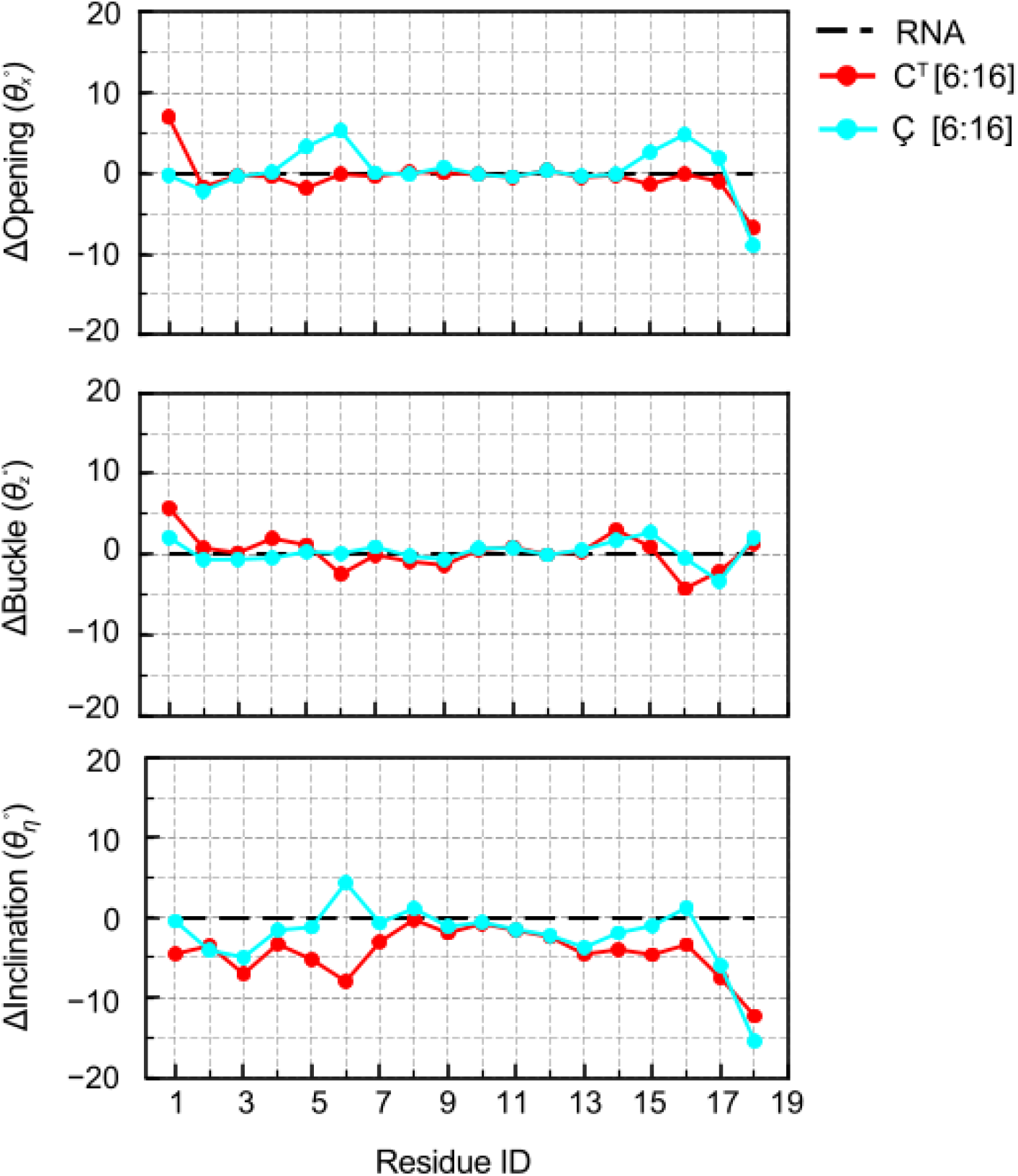
Comparison of change in average base pair parameters opening, buckle, and inclination for each residue from TEMPO and Ç labelled RNA duplexes with respect to unlabelled RNA duplex (∆= Labeled RNA base_(i)_ – Unlabelled RNA base_(i)_)

#### 3.3.2 Position based effect of N^T^

Differences in base pair geometry between the N^T^ labeled structures and the unlabeled RNA are seen only in the rotational parameters (Figure 3) and not in the translational parameters **(Supplementary Figure S4)**. Either minor differences or no differences are seen for opening, twist, tilt and tip **(Supplementary Figure S5)**. For C^T^[3:13] and C^T^[6:16] there is no opening of base pairs at the label attachment site and in A^T^[2:12] there is slight increase in opening (~5°) at the 12^th^ residue. In the G^T^[4:14] an opening (<5°) is observed in the base pairs succeeding the attachment sites. In A^T^[2:12] and C^T^[3:13] the perturbation in propeller parameter is extremely low in the interior of the duplex (i.e. the 12^th^ and the 13^th^ position) and at the termini (5’-end) there is an increase in perturbation in the base pairs preceding the attachment site. The same is not seen in G^T^[4:14] and C^T^[6:16]. In G^T^[4:14] there is perturbation only at the attachment site and in C^T^[6:16] there is an increase in propeller angle preceding the label attachment sites.

**Figure 3.**
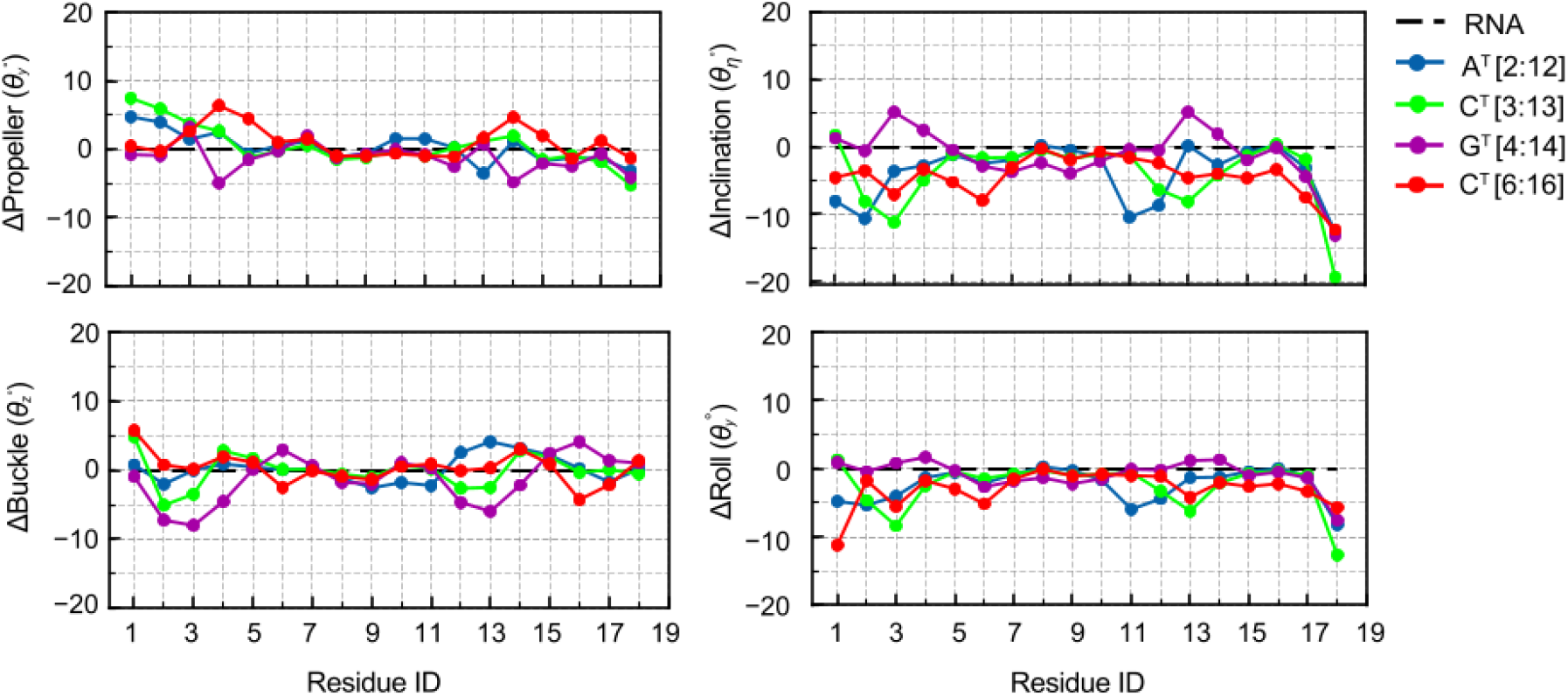
Comparison of change in average local base pair parameters for each residue from TEMPO labelled RNA duplexes at four attachment sites A^T^[2:12], C^T^[3:13], G^T^[4:14], and C^T^[6:16] with respect to unlabelled RNA duplex.

Buckling of <5° is observed in rest of the N^T^ labeled RNA duplexes and in G^T^[4:14], increased buckling of base pairs preceding to the attachment site is observed. Introduction of N^T^ at 2:12 and 3:13 and 6:16 results in change (≤8°) in the roll angles of base pairs preceding and succeeding the label attachment site. While G^T^[4:14], which protrudes out into the solvent, has positive inclination, the rest of N^T^ labels, which are in interior (**Supplementary Figure S6**), have negative inclination. Shear, stretch, and opening report on the hydrogen bond capabilities of a base pair.[39] The overall change in opening is relatively small and no changes in shear and stretch parameters are observed. This is in agreement with the percentage of hydrogen bonds (number of hydrogen bonds present in comparison to the total number of expected hydrogen bonds) between the two strands over the entire simulation (**Supplementary Table S1**), i.e. the change in local environment is not large enough to disturb the hydrogen bonding pattern. In summary, the effect of N^T^/Ç labels is restricted to the immediate vicinity of the labels and is dependent on the location of the label in the RNA.

### 3.4 Quadruplex RNA

Guanine-rich DNA/RNA sequences associate into G-quartet/tetrad stacks connected by short nucleotide loops and depending on experimental conditions, form a plethora of complex structures referred to as G-quadruplexes. These structures have been implicated in gene expression and regulation and mRNA biology and given their functional roles in cellular biology, there structural properties are of great interest.[40–43] In this study, the QRNA is parallel stranded propeller type G-quadruplex (**Figure 4**) with G-tetrads connected by “UUA” loops. Slight perturbations on the geometry of the base pair arrangements can be combined and associated to intrinsic flexibility of selected regions (or loops), affecting the distance distribution. Using the A^T^ spin probe, we can anticipate that slight changes of a single triad (“TTA/UUA” loop) of QRNA can be detected. Thus, the MD analysis has been performed on the unlabeled QRNA (QRNA_U_) and labeled QRNA structure (QRNA_L_) with TEMPO labels attached to adenines at 8^th^ position. These A^T^ labels are attached in the ‘UUA loop’ of the QRNA. For this system, we have studied the effect of A^T^ labels on the conformational stability and flexibility of the QRNA structure.

**Figure 4:**
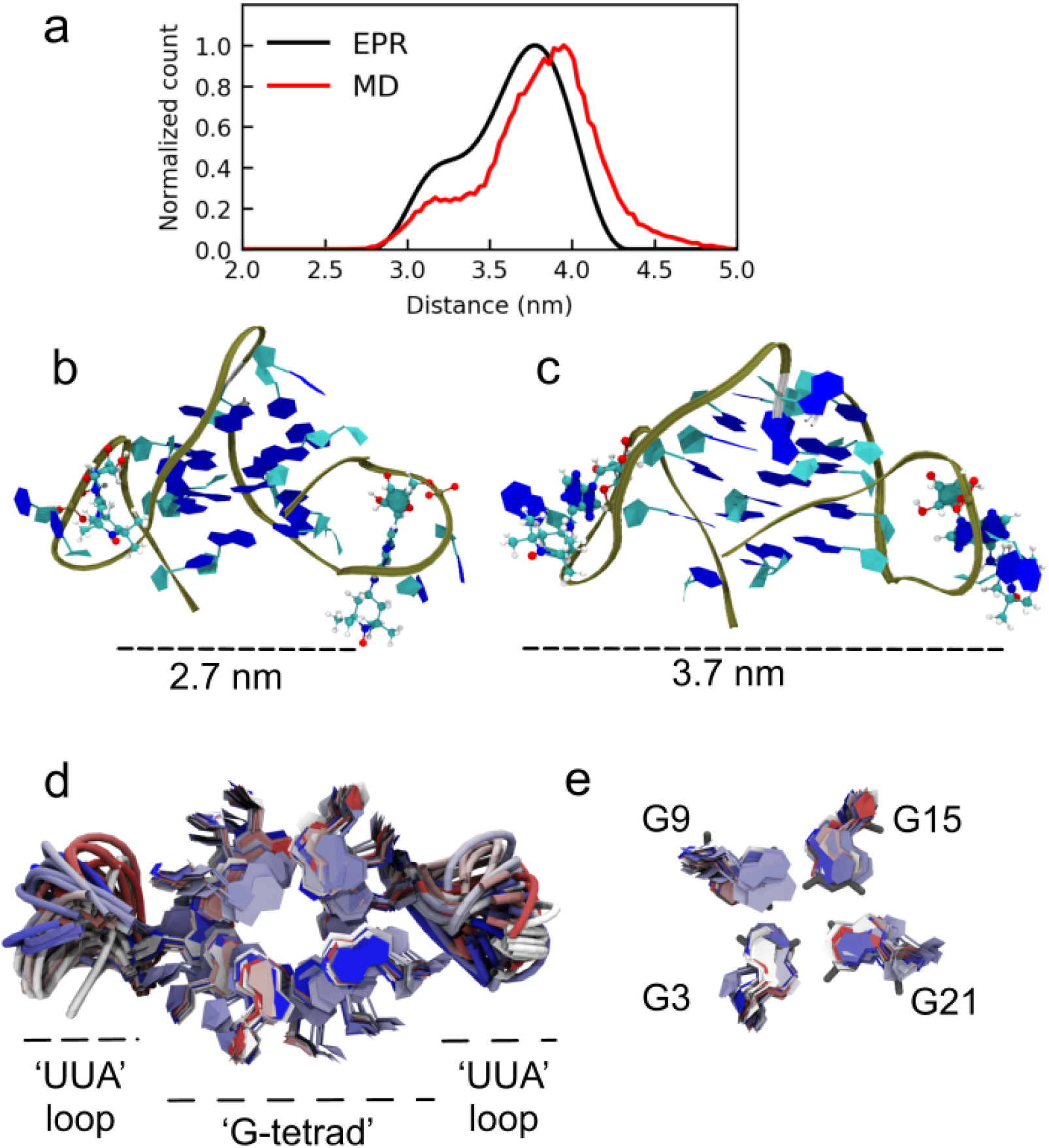
Quadruplex RNA conformational properties (**a**) Distribution of distance between the oxygen radicals of the two A^T^ labels attached to A8 and A20 in EPR experiments (black) and MD simulations (red) (**b & c**) Snapshots from QRNA MD ensemble showing the labelled QRNA conformations with distance between the two A^T^ labels at 2.7 nm and 3.7 nm. (d) Top view of the G-quadruplex showing conformations “UUA” loops and “G-quadruplex” core from the MD simulation ensemble taken at an interval of 1000 from an ensemble of 50,000 structures colored from a spectrum of blue-white-red. (e) Only the first G-quartet formed by G3, G9, G15, and G21. While “UUA” loops are flexible, G-tetrad core is rigid.

In our simulations, we observe high flexibility of the “UUA” loop both in labeled and unlabeled QRNA structure (**Figure 4d**). The “UUA” loops are highly flexible because there is no possibility of standard Watson-Crick base pairing as in a standard duplex. All atom RMSD reported by Islam et al.[44] for the G-quadruplex DNA (QDNA) was 0.45 nm34 which is very similar to the average all atom RMSD of QRNA_U_ (0.46 ± 0.05 nm) and QRNA_L_ (0.40 ± 0.05). Attachment of TEMPO labels at A8 does not affect the guanine core structure, which is observed in the calculated RMSD of the central guanine core for QRNA_U_ (0.27 nm) and QRNA_L_ (0.24 nm) simulations (**Figure 4e**). Our results are in agreement with the findings for the QDNA simulations,[44] i.e. “TTA/UUA” loops adopt a variety of conformations whereas the core guanine is rigid. G-quadruplex geometry was analyzed for both QRNA_U_ and QRNA_L_ and the difference in the geometry parameters is minor (**Table S2**). Stacking area, which is a representative measure of the strength of stacking interactions, is identical in both systems. Only a small difference of 0.9 °C is observed in the melting temperature between unlabeled and labeled QRNA samples (Table 1 in [7]), suggesting that TEMPO labels have a minimal or no effect on the QRNA structure.[7]

We monitored the r_OO_ distance between the two A^T^ labels attached to A8 in the QRNA_L_ structure and r_NL_ distance in the QRNA_U_ (**Table 2**). From the PELDOR experiments (see also reference [7], an asymmetric profile (negative skewness) has been observed (black curve) with two main conformations centred at 3.18 nm and 3.77 nm, respectively. A similar negatively skewed distribution is seen in MD simulations, with a shoulder around 3.25 nm and a peak at 4 nm in the QRNA_L_ data (**Figure 4a**). Analysis of the MD trajectories yielded a good agreement for the distance distribution (**Figure 4a**, **red curve**). The two distinct populations of structures present in the QRNA_L_ ensemble, arises from the diversity in conformations adopted by the “UUA” loops (**Figure 4b**), as seen by Islam et al.[44].

## 4. CONCLUSION

In the absence of X-ray crystallography or NMR based structural studies of such systems, these results provide valuable insights into the effects of nitroxide labels on the local geometry of nucleic acid molecules. The results of our molecular dynamics studies of unlabeled, N^T^ and Ç labeled RNA duplexes suggest that TEMPO or Ç labels do not affect the global structure at 298 K. The minor perturbations in the local environment of the labels, do not affect the hydrogen bonding capabilities of the nucleobases. Position based effect on the structure by N^T^ label is seen in our simulations suggesting that the point of attachment for the label should be chosen carefully. The design of new probes and even the pulsed EPR experiments will benefit of such detailed description of the geometry of modified base-pair. Further, as the synthesis of TEMPO-like spin probes is less demanding and since the effects of N^T^ and Ç are minor; the applicability of TEMPO-like spin probes can be extended without hesitations to RNA and DNA architectures of higher complexity.

## Supporting Information

Supplementary Figure S1 to S5, Tables S1 and S2, and the force field parameters of TEMPO and Ç labels.

## Notes

The authors declare no conflict of interest.

## ACKNOWLEDGEMENTS

SCD acknowledges the financial support of Max Planck Society and Volkswagen Foundation grant 83940. We are very grateful to Prof. Claudia Höbartner and Prof. Gerrit Groenhof for their valuable comments in structuring the final manuscript.

## References

[1] G. Jeschke, Dipolar Spectroscopy – Double-Resonance Methods, EMagRes. (2016) 1459–1476. https://doi.org/doi:10.1002/9780470034590.emrstm1518.

[2] G. Jeschke, The contribution of modern EPR to structural biology, Emerg. Top. Life Sci. 2 (2018) 9–18. https://doi.org/10.1042/ETLS20170143.

[3] A. Doll, G. Jeschke, Wideband frequency-swept excitation in pulsed EPR spectroscopy, J. Magn. Reson. 280 (2017) 46–62. https://doi.org/https://doi.org/10.1016/j.jmr.2017.01.004.

[4] T.H. Edwards, S. Stoll, Optimal Tikhonov regularization for DEER spectroscopy, J. Magn. Reson. 288 (2018) 58–68. https://doi.org/10.1016/j.jmr.2018.01.021.

[5] R. Dastvan, E.-M. Brouwer, D. Schuetz, O. Mirus, E. Schleiff, T.F. Prisner, Relative Orientation of POTRA Domains from Cyanobacterial Omp85 Studied by Pulsed EPR Spectroscopy, Biophys. J. 110 (2016) 2195–2206. https://doi.org/10.1016/j.bpj.2016.04.030.

[6] L.S. Stelzl, N. Erlenbach, M. Heinz, T.F. Prisner, G. Hummer, Resolving the Conformational Dynamics of DNA with Ångstrom Resolution by Pulsed Electron– Electron Double Resonance and Molecular Dynamics, J. Am. Chem. Soc. 139 (2017) 11674–11677. https://doi.org/10.1021/jacs.7b05363.

[7] G. Sicoli, F. Wachowius, M. Bennati, C. Höbartner, Probing secondary structures of spin-labeled RNA by pulsed EPR spectroscopy., Angew. Chem. Int. Ed. Engl. 49 (2010) 6443–7. https://doi.org/10.1002/anie.201000713.

[8] J.L. Sarver, J.E. Townsend, G. Rajapakse, L. Jen-Jacobson, S. Saxena, Simulating the dynamics and orientations of spin-labeled side chains in a protein-DNA complex., J. Phys. Chem. B. 116 (2012) 4024–33. https://doi.org/10.1021/jp211094n.

[9] O. Duss, E. Michel, M. Yulikov, M. Schubert, G. Jeschke, F.H.-T. Allain, Structural basis of the non-coding RNA RsmZ acting as a protein sponge, Nature. 509 (2014) 588–592. https://doi.org/10.1038/nature13271.

[10] Z. Wu, A. Feintuch, A. Collauto, L.A. Adams, L. Aurelio, B. Graham, G. Otting, D. Goldfarb, Selective Distance Measurements Using Triple Spin Labeling with Gd3+, Mn2+, and a Nitroxide, J. Phys. Chem. Lett. 8 (2017) 5277–5282. https://doi.org/10.1021/acs.jpclett.7b01739.

[11] E.R. Georgieva, A.S. Roy, V.M. Grigoryants, P.P. Borbat, K.A. Earle, C.P. Scholes, J.H. Freed, Effect of freezing conditions on distances and their distributions derived from Double Electron Electron Resonance (DEER): A study of doubly-spin-labeled T4 lysozyme, J. Magn. Reson. 216 (2012) 69–77. https://doi.org/https://doi.org/10.1016/j.jmr.2012.01.004.

[12] K. Halbmair, J. Seikowski, I. Tkach, C. Höbartner, D. Sezer, M. Bennati, High-resolution measurement of long-range distances in RNA: pulse EPR spectroscopy with TEMPO-labeled nucleotides, Chem. Sci. 7 (2016) 3172–3180. https://doi.org/10.1039/C5SC04631A.

[13] C. Prior, L. Danilāne, V.S. Oganesyan, All-atom molecular dynamics simulations of spin labelled double and single-strand DNA for EPR studies, Phys. Chem. Chem. Phys. 20 (2018) 13461–13472. https://doi.org/10.1039/C7CP08625C.

[14] G. Sicoli, G. Mathis, S. Aci-Sèche, C. Saint-Pierre, Y. Boulard, D. Gasparutto, S. Gambarelli, Lesion-induced DNA weak structural changes detected by pulsed EPR spectroscopy combined with site-directed spin labelling, Nucleic Acids Res. 37 (2009) 3165–3176. https://doi.org/10.1093/nar/gkp165.

[15] M. Heinz, N. Erlenbach, L.S. Stelzl, G. Thierolf, N.R. Kamble, S.T. Sigurdsson, T.F. Prisner, G. Hummer, High-resolution EPR distance measurements on RNA and DNA with the non-covalent ◻ spin label, Nucleic Acids Res. 48 (2020) 924–933. https://doi.org/10.1093/nar/gkz1096.

[16] O. Schiemann, P. Cekan, D. Margraf, T.F. Prisner, S.T. Sigurdsson, Relative orientation of rigid nitroxides by PELDOR: beyond distance measurements in nucleic acids., Angew. Chem. Int. Ed. Engl. 48 (2009) 3292–5. https://doi.org/10.1002/anie.200805152.

[17] H. Martadinata, A.T. Phan, Structure of Propeller-Type Parallel-Stranded RNA G-Quadruplexes, Formed by Human Telomeric RNA Sequences in K+ Solution, J. Am. Chem. Soc. 131 (2009) 2570–2578. https://doi.org/10.1021/ja806592z.

[18] S. Pronk, S. Páll, R. Schulz, P. Larsson, P. Bjelkmar, R. Apostolov, M.R. Shirts, J.C. Smith, P.M. Kasson, D. Van Der Spoel, B. Hess, E. Lindahl, GROMACS 4.5: A high-throughput and highly parallel open source molecular simulation toolkit, Bioinformatics. 29 (2013). https://doi.org/10.1093/bioinformatics/btt055.

[19] A. Pérez, I. Marchán, D. Svozil, J. Sponer, T.E. Cheatham, C. a Laughton, M. Orozco, Refinement of the AMBER force field for nucleic acids: improving the description of alpha/gamma conformers., Biophys. J. 92 (2007) 3817–29. https://doi.org/10.1529/biophysj.106.097782.

[20] E. Stendardo, A. Pedone, P. Cimino, M. Cristina Menziani, O. Crescenzi, V. Barone, Extension of the AMBER force-field for the study of large nitroxides in condensed phases: an ab initio parameterization., Phys. Chem. Chem. Phys. 12 (2010) 11697–709. https://doi.org/10.1039/c001481h.

[21] C.I. Bayly, P. Cieplak, W. Cornell, P.A. Kollman, A well-behaved electrostatic potential based method using charge restraints for deriving atomic charges: the RESP model, J. Phys. Chem. 97 (1993) 10269–10280. https://doi.org/10.1021/j100142a004.

[22] P. Cieplak, W.D. Cornell, C. Bayly, P.A. Kollman, Multiconformational RESP Methodology to Biopolymers◻: Charge Derivation for, 16 (1995).

[23] H.J.C. Berendsen, J.P.M. Postma, W.F. van Gunsteren, a. DiNola, J.R. Haak, Molecular dynamics with coupling to an external bath, J. Chem. Phys. 81 (1984) 3684. https://doi.org/10.1063/1.448118.

[24] S. Nosé, A molecular dynamics method for simulations in the canonical ensemble, Mol. Phys. 52 (1984) 255–268. https://doi.org/10.1080/00268978400101201.

[25] W.G. Hoover, Canonical dynamics: Equilibrium phase-space distributions, Phys. Rev. A. 31 (1985) 1695–1697. https://doi.org/10.1103/PhysRevA.31.1695.

[26] M. Parrinello, A. Rahman, Polymorphic transitions in single crystals: A new molecular dynamics method, J. Appl. Phys. 52 (1981) 7182–7190. https://doi.org/10.1063/1.328693.

[27] B. Hess, H. Bekker, H.J.C. Berendsen, J.G.E.M. Fraaije, LINCS: A linear constraint solver for molecular simulations, J. Comput. Chem. 18 (1997) 1463–1472. https://doi.org/10.1002/(SICI)1096-987X(199709)18:12<1463::AID-JCC4>3.0.CO;2-H.

[28] T. Darden, D. York, L. Pedersen, Particle mesh Ewald: An N⋅log(N) method for Ewald sums in large systems, J. Chem. Phys. 98 (1993) 10089–10092. https://doi.org/10.1063/1.464397.

[29] U. Essmann, L. Perera, M.L. Berkowitz, T. Darden, H. Lee, L.G. Pedersen, A smooth particle mesh Ewald method, J. Chem. Phys. 103 (1995) 8577–8593. https://doi.org/10.1063/1.470117.

[30] C. Höbartner, G. Sicoli, F. Wachowius, D.B. Gophane, S.T. Sigurdsson, Synthesis and characterization of RNA containing a rigid and nonperturbing cytidine-derived spin label., J. Org. Chem. 77 (2012) 7749–54. https://doi.org/10.1021/jo301227w.

[31] I. Tkach, S. Pornsuwan, C. Höbartner, F. Wachowius, S.T. Sigurdsson, T.Y. Baranova, U. Diederichsen, G. Sicoli, M. Bennati, Orientation selection in distance measurements between nitroxide spin labels at 94 GHz EPR with variable dual frequency irradiation, Phys. Chem. Chem. Phys. 15 (2013) 3433–3437. https://doi.org/10.1039/C3CP44415E.

[32] R. Kumar, H. Grubmüller, do_x3dna: a tool to analyze structural fluctuations of dsDNA or dsRNA from molecular dynamics simulations, Bioinformatics. 31 (2015) 2583–2585. https://doi.org/10.1093/bioinformatics/btv190.

[33] X.-J. Lu, H.J. Bussemaker, W.K. Olson, DSSR: an integrated software tool for dissecting the spatial structure of RNA, Nucleic Acids Res. 43 (2015) e142–e142. https://doi.org/10.1093/nar/gkv716.

[34] L. Schrodinger, The PyMOL Molecular Graphics System Version 1.7, (2015).

[35] W. Humphrey, A. Dalke, K. Schulten, VMD: Visual molecular dynamics, J. Mol. Graph. 14 (1996) 33–38. https://doi.org/10.1016/0263-7855(96)00018-5.

[36] P. Auffinger, E. Westhof, Hydrophobic Groups Stabilize the Hydration Shell of 2′-O-Methylated RNA Duplexes, Angew. Chemie Int. Ed. 40 (2001) 4648–4650. https://doi.org/10.1002/1521-3773(20011217)40:24<4648::AID-ANIE4648>3.0.CO;2-U.

[37] M. Popenda, E. Biala, J. Milecki, R.W. Adamiak, Solution structure of RNA duplexes containing alternating CG base pairs: NMR study of r(CGCGCG) 2 and 2′-O - Me(CGCGCG) 2 under low salt conditions, Nucleic Acids Res. 25 (1997) 4589–4598. https://doi.org/10.1093/nar/25.22.4589.

[38] R.E. Dickerson, Definitions and nomenclature of nucleic acid structure components, Nucleic Acids Res. 17 (1989) 1797–1803. https://doi.org/10.1093/nar/17.5.1797.

[39] X.-J. Lu, W.K. Olson, 3DNA: a versatile, integrated software system for the analysis, rebuilding and visualization of three-dimensional nucleic-acid structures., Nat. Protoc. 3 (2008) 1213–27. https://doi.org/10.1038/nprot.2008.104.

[40] M.M. Fay, S.M. Lyons, P. Ivanov, RNA G-Quadruplexes in Biology: Principles and Molecular Mechanisms, J. Mol. Biol. 429 (2017) 2127–2147. https://doi.org/10.1016/j.jmb.2017.05.017.

[41] R. Hänsel-Hertsch, M. Di Antonio, S. Balasubramanian, DNA G-quadruplexes in the human genome: detection, functions and therapeutic potential, Nat. Rev. Mol. Cell Biol. 18 (2017) 279–284. https://doi.org/10.1038/nrm.2017.3.

[42] C.K. Kwok, C.J. Merrick, G-Quadruplexes: Prediction, Characterization, and Biological Application, Trends Biotechnol. 35 (2017) 997–1013. https://doi.org/10.1016/j.tibtech.2017.06.012.

[43] S. Millevoi, H. Moine, S. Vagner, G-quadruplexes in RNA biology, WIREs RNA. 3 (2012) 495–507. https://doi.org/10.1002/wrna.1113.

[44] B. Islam, M. Sgobba, C. Laughton, M. Orozco, J. Sponer, S. Neidle, S. Haider, Conformational dynamics of the human propeller telomeric DNA quadruplex on a microsecond time scale, Nucleic Acids Res. 41 (2013) 2723–2735. https://doi.org/10.1093/nar/gks1331.

