## Supplementary Material for "The ‘hidden side’ of spin labeled oligonucleotides: Molecular Dynamics study focusing on the EPR-silent components of base pairing"

**Supplementary figures**

**Supplementary Figure S1:** Distance distribution obtained by (a) Tikhonov regularization (b) Gaussian-fit.^[1]^

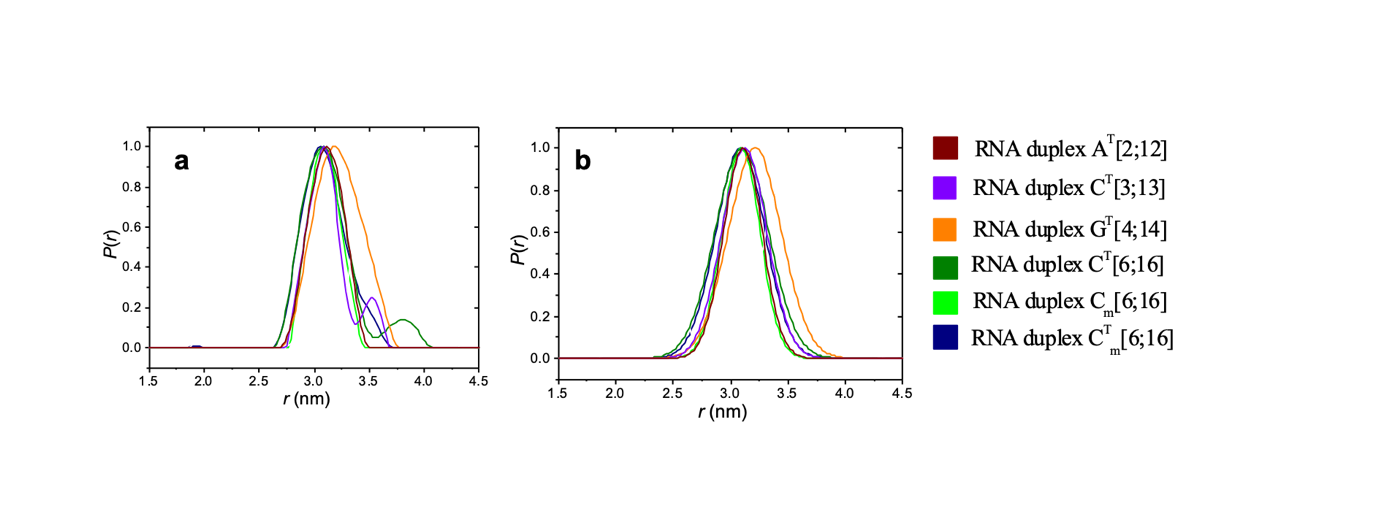

**Supplementary Figure S2:** Comparison of change in average base pair parameters between C^T^[6:16] and Ç [6:16] with respect to unlabelled RNA. There is no change in these parameters with respect to unlabelled RNA.

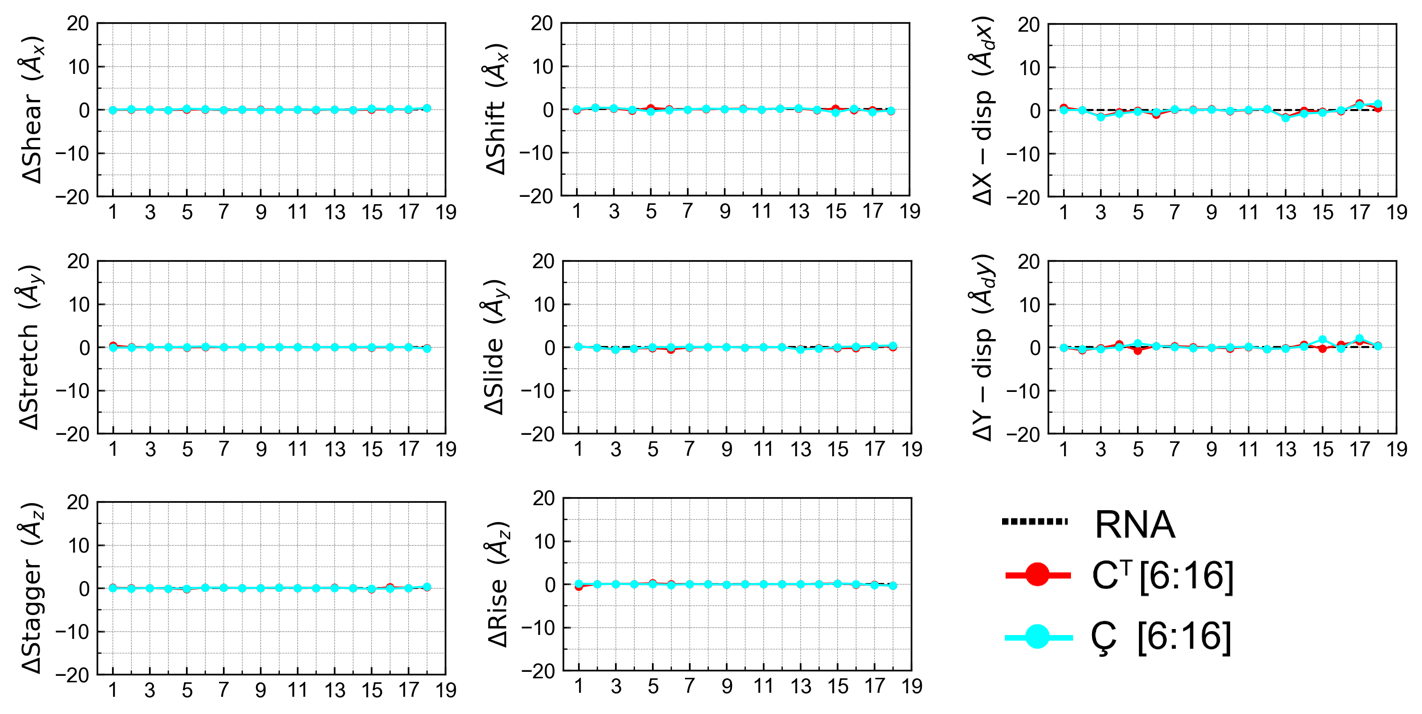

**Supplementary Figure S3:** Comparison of change in average base pair parameters between C^T^[6:16] and Ç[6:16] with respect to unlabelled RNA. Twist and Tip profiles are identical between the two labels and increased rolling and propeller are seen for C^T^ labelled structure compared to Ç.

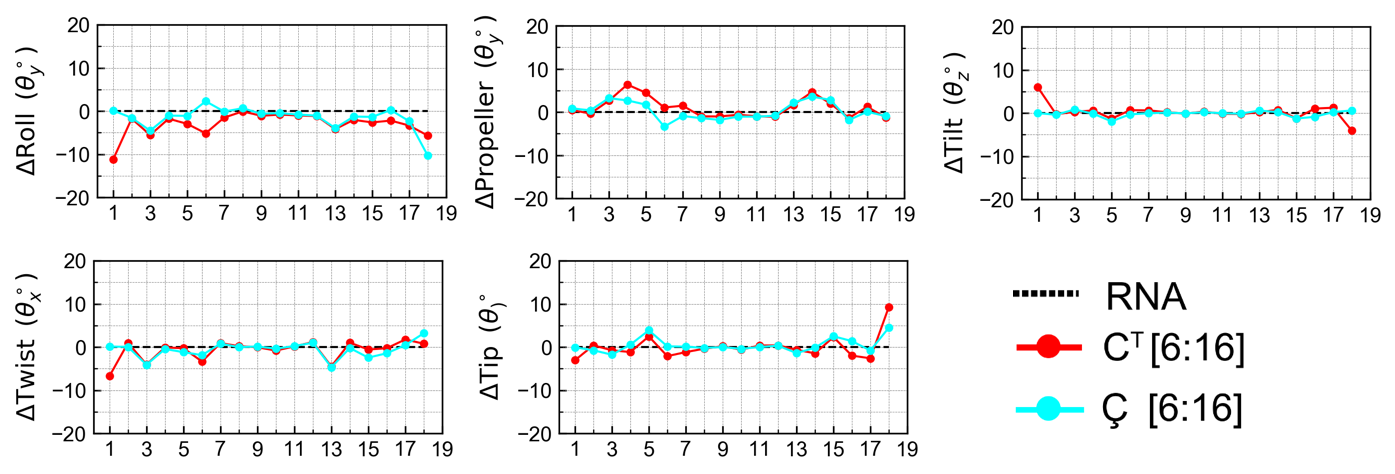

**Supplementary Figure S4:** Comparison of change in average base pair parameter for each residue from N^T^ labeled RNA duplex with respect to unlabeled RNA duplex.

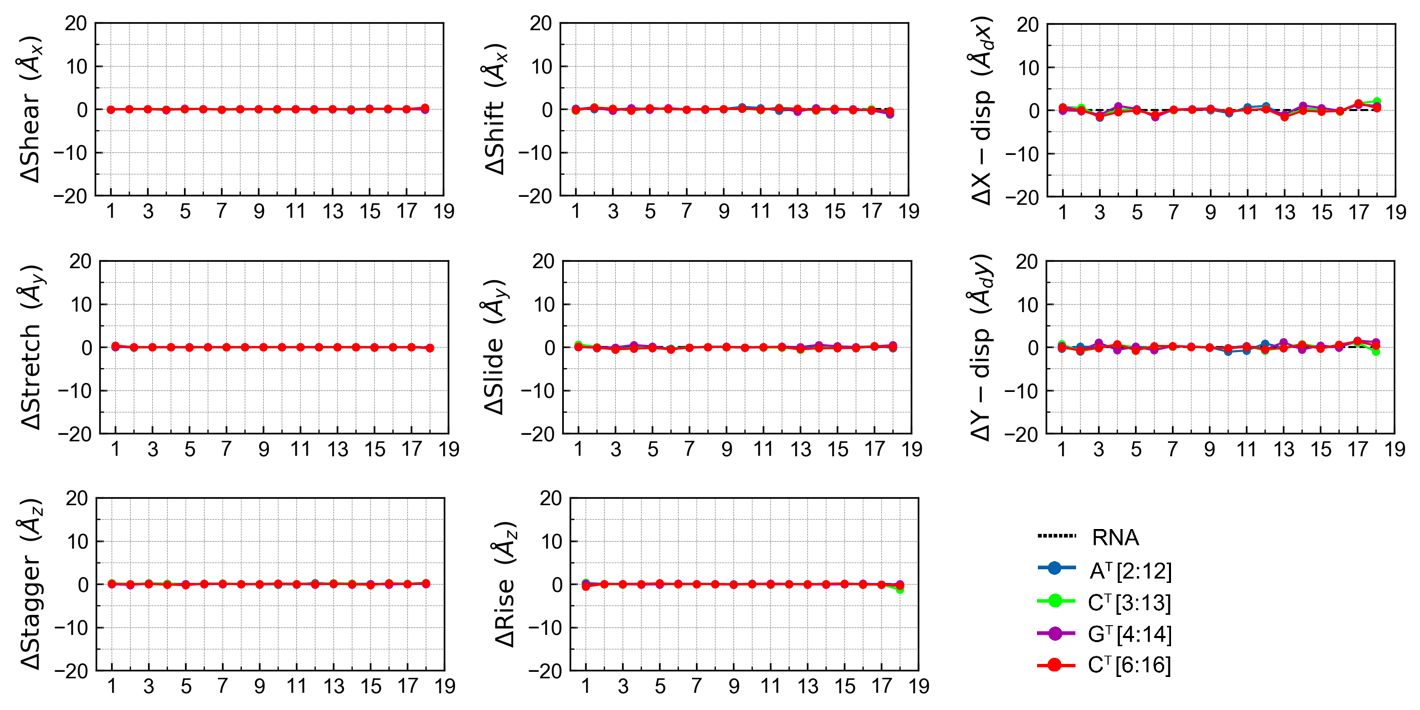

**Supplementary Figure S5:** Comparison of change in average base pair parameter for each residue from TEMPO labeled RNA duplex with respect to unllabeled RNA duplex.

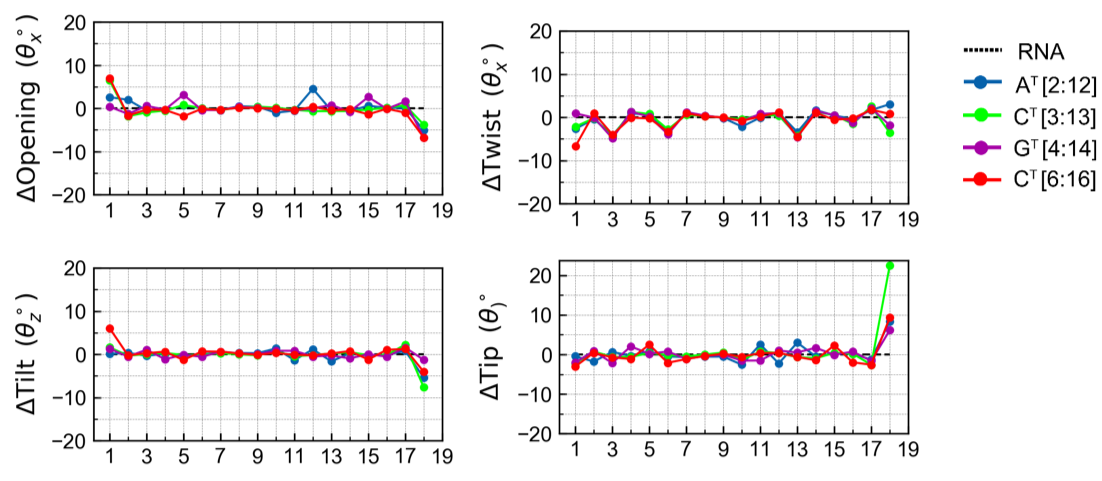

**Supplementary Figure S6:** Orientation and position of TEMPO labels A^T^ (blue: attached at A2:A12), C^T^ (green: attached at C3:C13), G^T^ (purple: attached at G4:G14), and C^T^ (red: attached at C6:C16), a) side view b) top view

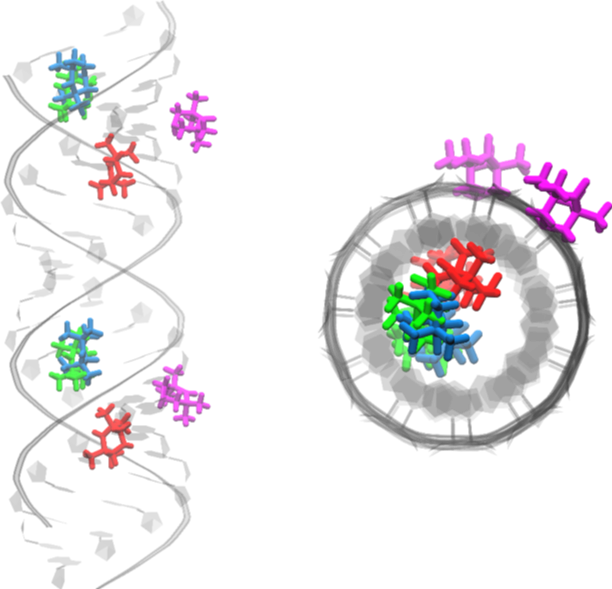

**Supplementary tables**

**Table S1:** Percentage of hydrogen bonds is calculated as the total number of expected hydrogen bonds between the two strands present during the entire length of the simulation in the labeled and unlabeled RNA molecules. The first two residues and the last two residues were not included in this analysis, since we observe opening of base pairs present at the termini.

| Simulation label | Percentage of hydrogen bonds  (±standard deviation) |
| --- | --- |
| RNA | 97±1.8 |
| RNA_m_[6:16] | 98±1.5 |
| A^T^[2:12] | 98±1.2 |
| C^T^[3:13] | 98±1.2 |
| G^T^[4:14] | 98±1.4 |
| C^T^[6:16] | 99±1.4 |
| C^T^_m_[6:16] | 99±1.3 |
| Ç[6:16] | 98±1.4 |
| Ç_m_[6:16] | 98±1.4 |

C_m_/C^T^_m_: 2’-OH of cytosine is methylated.

**Table S2:** Comparison of G-quadraplex geometry parameters, between the unlabeled quadraplex-RNA (QRNA_U_) and A^T^ labeled quadraplex-RNA (QRNA_L_) at 8^th^ nucleotide, for the first two G-tetrads out of the three.

| Simulation label | G-tetrad number | Stacking area  (±standard deviation) (nm^2^) | Rise (nm) | Twist (º) |
| --- | --- | --- | --- | --- |
| QRNA_U_ | 1 | 0.15±0.03 | 0.32±0.04 | 26.69±2.75 |
|  | 2 | 0.12±0.02 | 0.40±0.03 | 26.76±2.27 |
| QRNA_L_ | 1 | 0.13±0.02 | 0.35±0.04 | 30.02±2.66 |
|  | 2 | 0.14±0.02 | 0.39±0.03 | 25.63±1.94 |

**Forcefield parameters in GROMACS format**

; Methylated Cytosine

[ MRC ]

[ atoms ]

P	P	1.08784	1

O1P	O2	-0.76666	2

O2P	O2	-0.76666	3

O5'	OS	-0.47250	4

C5'	CT	0.12894	5

H5'1	H1	0.04256	6

H5'2	H1	0.04256	7

C4'	CT	0.15216	8

O4'	OS	-0.46523	9

C1'	CT	0.36862	10

C2'	CT	0.04045	11

O2'	OS	-0.32773	12

CM2	CT	-0.03845	13

HM'1	H1	0.06505	14

HM'2	H1	0.06505	15

HM'3	H1	0.06505	16

H2'	H1	0.09036	17

N1	N*	-0.21522	18

C2	C	0.88670	19

O2	O	-0.65599	20

N3	NC	-0.81275	21

C4	CA	0.90203	22

N4	N2	-0.99189	23

H41	H	0.42507	24

H42	H	0.42507	25

C5	CM	-0.59721	26

C6	CM	0.12624	27

H6	H4	0.18750	28

H5	HA	0.20229	29

H1'	H2	0.04166	30

H4'	H1	0.03942	31

C3'	CT	0.06749	32

H3'	H1	0.14600	33

O3'	OS	-0.48782	34

; Unmethylated Ç

[ RCM ]

[ atoms ]

P	P	1.16620	1

O1P	O2	-0.77600	2

O2P	O2	-0.77600	3

O5'	OS	-0.49890	4

C5'	CI	0.05580	5

H5'1	H1	0.06790	6

H5'2	H1	0.06790	7

C4'	CT	0.10650	8

H4'	H1	0.11740	9

O4'	OS	-0.35480	10

C1'	CT	0.00220	11

H1'	H2	0.22618	12

N1	N*	-0.18852	13

C6	CM	-0.10657	14

H6	H4	0.25271	15

C5	CCT	0.03991	16

C4	CCT	0.64239	17

N4	NAT	-0.49581	18

H41	HNT	0.36906	19

C21	CAT	0.10457	20

C3	CAT	0.41424	21

C1	CAT	-0.30511	22

H1	HAT	0.20632	23

O21	OST	-0.39500	24

C9	CAT	-0.04431	25

C10	CAT	-0.13765	26

C11	CT	0.48424	27

C41	CAT	-0.43017	28

H2	HAT	0.22330	29

C13	CT	-0.43608	30

H5	HN	0.11556	31

H61	HN	0.11556	32

H7	HN	0.11556	33

C14	CT	-0.43608	34

H8	HN	0.11556	35

H9	HN	0.11556	36

H10	HN	0.11556	37

N7	NN	0.03466	38

O3	ON	-0.17157	39

LP1	LP	-0.11000	40

LP2	LP	-0.11000	41

C12	CT	0.52037	42

C15	CT	-0.43687	43

H11	HN	0.11220	44

H12	HN	0.11220	45

H13	HN	0.11220	46

C16	CT	-0.43687	47

H14	HN	0.11220	48

H15	HN	0.11220	49

H16	HN	0.11220	50

N3	NC	-0.82919	51

C2	C	0.94499	52

O2	O	-0.64590	53

C3'	CT	0.20220	54

H3'	H1	0.06150	55

C2'	CT	0.06700	56

H2'1	H1	0.09720	57

O2'	OH	-0.61390	58

HO'2	HO	0.41860	59

O3'	OS	-0.52460	60

; Ç

[ MCM ]

[ atoms ]

P	P	1.16620	1

O1P	O2	-0.77600	2

O2P	O2	-0.77600	3

O5'	OS	-0.49890	4

C5'	CI	0.05580	5

H5'1	H1	0.06790	6

H5'2	H1	0.06790	7

C4'	CT	0.10650	8

H4'	H1	0.11740	9

O4'	OS	-0.35480	10

C1'	CT	0.00220	11

H1'	H2	0.22618	12

N1	N*	-0.18852	13

C6	CM	-0.10657	14

H6	H4	0.25271	15

C5	CCT	0.03991	16

C4	CCT	0.64239	17

N4	NAT	-0.49581	18

H41	HNT	0.36906	19

C21	CAT	0.10457	20

C3	CAT	0.41424	21

C1	CAT	-0.30511	22

H1	HAT	0.20632	23

O21	OST	-0.39500	24

C9	CAT	-0.04431	25

C10	CAT	-0.13765	26

C11	CT	0.48424	27

C41	CAT	-0.43017	28

H2	HAT	0.22330	29

C13	CT	-0.43608	30

H5	HN	0.11556	31

H61	HN	0.11556	32

H7	HN	0.11556	33

C14	CT	-0.43608	34

H8	HN	0.11556	35

H9	HN	0.11556	36

H10	HN	0.11556	37

N41	NN	0.03466	38

O3	ON	-0.17157	39

LP1	LP	-0.11000	40

LP2	LP	-0.11000	41

C12	CT	0.52037	42

C15	CT	-0.43687	43

H11	HN	0.11220	44

H12	HN	0.11220	45

H13	HN	0.11220	46

C16	CT	-0.43687	47

H14	HN	0.11220	48

H15	HN	0.11220	49

H16	HN	0.11220	50

N3	NC	-0.82919	51

C2	C	0.94499	52

O2	O	-0.64590	53

C3'	CT	0.20220	54

H3'	H1	0.06150	55

C2'	CT	0.06700	56

H2'1	H1	0.09720	57

O2'	OS	-0.61390	58

C22	CT	0.08860	59

H25	H1	0.11000	60

H26	H1	0.11000	61

H27	H1	0.11000	62

O3'	OS	-0.52460	63

**;Methylated Cytosine attached to TEMPO (C^T^)**

[ MCN ]

P	P	1.16620	1

O1P	O2	-0.77600	2

O2P	O2	-0.77600	3

O5'	OS	-0.49890	4

C5'	CT	0.05580	5

H5'1	H1	0.06790	6

H5'2	H1	0.06790	7

C4'	CT	0.10650	8

H4'	H1	0.11740	9

O4'	OS	-0.35480	10

C1'	CT	0.00660	11

H1'	H2	0.20290	12

N1	N*	-0.04840	13

C6	CM	0.00530	14

H6	H4	0.19580	15

C5	CM	-0.52150	16

H5	HA	0.19280	17

C4	CA	0.81850	18

NL	N2	-0.76788	19

HNL	H	0.35583	20

N3	NC	-0.75840	21

C2	C	0.75380	22

O2	O	-0.62520	23

C3'	CT	0.20220	24

H3'	H1	0.06150	25

C2'	CT	0.06700	26

H2'1	H1	0.09720	27

O2'	OS	-0.61390	28

C22	CT	0.08860	29

H25	H1	0.11000	30

H26	H1	0.11000	31

H27	H1	0.11000	32

O3'	OS	-0.52460	33
